# Subtype-specific epigenomic landscape and 3D genome structure in bladder cancer

**DOI:** 10.1101/2020.02.26.966697

**Authors:** Tejaswi Iyyanki, Baozhen Zhang, Qiushi Jin, Hongbo Yang, Tingting Liu, Xiaotao Wang, Jie Xu, Fan Song, Yu Luan, Hironobu Yamashita, Lu Wang, Joshua Warrick, Jay Raman, Joshua J. Meeks, David Degraff, Feng Yue

**Author notes:** These authors contributed equally to this work (Tejaswi Iyyanki, Baozhen Zhang).

## Abstract

Muscle-invasive bladder cancers have recently been characterized by their distinct expression of luminal and basal genes, which could be used to predict key clinical features such as disease progression and overall survival. For example, FOXA1, GATA3, and PPARG have been shown to be essential for luminal subtype-specific regulation and subtype switching, while TP63 and STAT3 are critical for basal subtype bladder cancer. Despite these advances, the underlying epigenetic mechanism and 3D chromatin architecture for subtype-specific regulation in bladder cancers remains largely unknown. Here, we determined the genome-wide transcriptome, enhancer landscape, TF binding profiles (FOXA1 and GATA3) in luminal and basal subtypes of bladder cancers. Furthermore, we mapped genome-wide chromatin interactions by Hi-C in both bladder cancer cell lines and primary patient tumors, for the first time in bladder cancer. We showed that subtype-specific transcription is accompanied by specific open chromatin and epigenomic marks, at least partially driven by distinct TF binding at distal-enhancers of luminal and basal bladder cancers. Finally, we identified a novel clinically relevant transcriptional factor, Neuronal PAS Domain Protein 2 (NPAS2), in luminal bladder cancers that regulates other luminal-specific genes (such as FOXA1, GATA3, and PPARG) and affects cancer cell proliferation and migration. In summary, our work shows a subtype-specific epigenomic and 3D genome structure in urinary bladder cancers and suggested a novel link between the circadian TF NPAS2 and a clinical bladder cancer subtype.

## Introduction

Urinary bladder cancers (BLCA) are the second most commonly diagnosed cancer in the United States, with over 81,400 total new cases diagnosed in 2019 [1, 2]. As BLCA is a morbid disease that is costly to treat, increased molecular understanding is required [3]. Expression of luminal (TFs – FOXA1, GATA3, PPARG, etc.) and basal (higher molecular weight keratins – KRT1, KRT5, KRT6A, etc.) [4, 5] genes have been used to molecularly characterize muscle invasive BLCA. In particular, the presence of basal BLCA, which is often enriched for squamous differentiation is associated with significant morbidity, disease progression and lower survival [6, 7].

Recent studies have identified both luminal (FOXA1, GATA3 and PPARG [8]) and basal (TP63 [9-12], STAT3 [4, 13-15], TFAP2A and TFAP2C [16]) -specific transcription factors (TFs) with functional roles in BLCA. For instance, we previously reported that FOXA1 and GATA3 cooperates with PPARG activation to drive trans-differentiation of basal BLCA cells to luminal subtype [8]. This observation is in agreement with the role of these factors in maintaining urothelial cell differentiation [17, 18] and suggests that inactivation of FOXA1, GATA3 and/or PPARG may be required for the emergence of a basal gene expression pattern during tumor progression. Subtype-specific TFs also appear to drive programs in oncogenesis and tumor progression. For example, basal factors STAT3, TP63 and TFAP2A/C have shown to promote tumor cell invasion and/or metastasis [12, 13, 16]. Similarly loss of FOXA1 [19] and GATA3 [20] have been shown to promote cell migration and invasion. Therefore, specific TFs play a key role in subtype specification.

In addition to directly regulating transcription, studies show TFs regulate gene expression through epigenetic histone modifications and open chromatin accessibility in breast cancers [21-24]. However, a comprehensive understanding of how TF repertoire, epigenetic open chromatin TF accessibility, histone modifications and 3D genome architecture that allows for subtype expression and hence its distinction is not well understood. Therefore, we performed the most comprehensive set of genome-wide experiments to systematically map the epigenome, transcriptome, TF binding and 3D chromatin contacts. *To our knowledge this is the first report identifying 3D genome architecture in bladder cancer and also use it to distinguish its subtypes*. Our work highlights the relevance of epigenetic modifications, open chromatin accessibility, TF repertoire and identifies a novel identified basic helix loop helix (bHLH) TF NPAS2 to coordinate the regulation of gene expression in bladder cancers.

## Results

### Comprehensive epigenomic profiling in both BLCA lines and primary tumors

In this project, we performed RNA-Seq, ChIP-Seq for Histone 3 lysine 27 acetylation (H3K27ac), Assay for Transposase-Accessible Chromatin using sequencing (ATAC-Seq) and genome-wide chromatin confirmation capture experiments (Hi-C) on 4 bladder cancer cell lines (Fig. 1A and 1B), two of which (RT4 and SW780) are previously annotated as luminal and the other two (SCABER and HT1376) characterized as basal subtype based on gene expression profiling [8, 25]. More importantly, we performed an identical set of experiments on two patient muscle-invasive bladder tumors as well (Fig 1B). Each experiment has at least two biological replicates (Supplementary table S1) and observed high correlation between the two replicates (Supplementary table S2). To our knowledge, the is the most comprehensive epigenomic profiling and the first to describe genome-wide 3D genome architecture of muscle-invasive BLCA.

**Figure 1:**
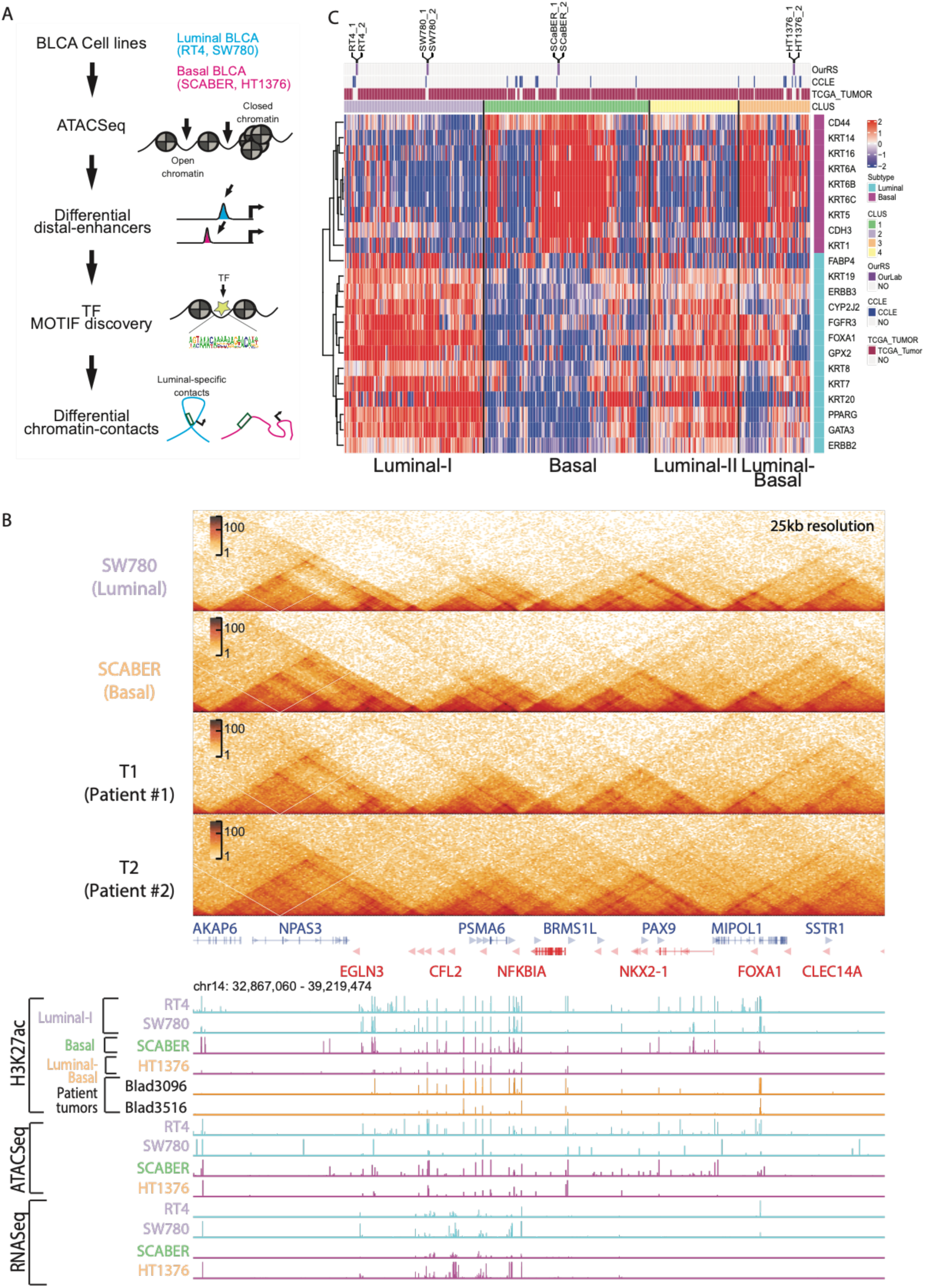
Comprehensive of epigenomic profiling in both bladder cancer lines and primary tumors. A) Schematic workflow of our study is shown here. We selected four muscle-invasive BLCA cell lines with known luminal (RT4 and SW780) and basal (SCABER and HT1376) subtypes and performed ATAC-Seq to identify open chromatins at regulatory elements, H3K27ac ChIP-Seq to define regulatory enhancers and performed TF motif discovery to identify luminal and basal-squamous specific TFs. We then performed Hi-C analysis to identify subtype-specific chromatin contacts. B) Genome browser signal tracks from an area of chromosome 14 near FOXA1 gene following comprehensive epigenomic analysis (H3K27ac, ATAC-seq, and RNA-seq) of RT4, SW780, SCaBER, HT1376 BLCA cell lines, as well as H3K27ac data from two primary BLCA specimens are shown. C) Hierarchical clustering of top 5,000 most variable genes in RNASeq data shows that there are four main clusters of molecular subtype where the four BLCA cell lines cluster amongst TCGA patients (top bar “TCGA_Tumor”) and CCLE bladder cancer cell lines (shown as a top bar “CCLE”) with distinct expression of luminal and basal genes..

### Transcriptome analysis of luminal and basal subtypes of muscle-invasive BLCA

To confirm the subtype classification of our selected cell lines, we first compared our RNA-Seq data from the 4 cell lines with the RNA-Seq data of 25 bladder cancer cell lines from the Cancer Cell Line Encyclopedia (CCLE [26]) and the RNA-Seq data of 408 patient bladder tumors from The Cancer Genome Atlas (TCGA [27]). First, we observed four major subtypes (Luminal I, Luminal II, Basal, Luminal and Basal) in our clustering analysis with both CCLE and TCGA data (Fig. S1A and S1B), consistent with previous report [7]. Second, we confirmed that our cell lines RT4 and SW780 were clustered with other luminal subtypes with similar gene signature (*FOXA1, KRT19, ERBB3, CYP2J2, GATA3, PPARG*, etc). Transcriptional data from our cultured SCABER cells clustered with other basal cell lines and HT1376 is clustered with type 4 (Basal/Luminal) (Fig. 1C). Finally, we performed Gene Ontology (GO) analysis for each groups and observed that while both Luminal-I and Luminal-II genes are enriched for genes important for NABA matrisome associated functions, Luminal I is more enriched for tumor microenvironment pathways such as leukocyte migration, cytokine production and blood vessel development (Fig. S1C) and Luminal II is enriched for transport processes, epithelial differentiation and skeletal muscle development (Fig. S1D). Differential gene expression analysis (DEG) (Fig. 2A) between Luminal I and Basal clusters revealed 2,753 upregulated basal genes involved in tumor microenvironment GO terms such as extracellular matrix organization, leukocyte activation, cytokine production and pathway and immune reactions and response pathways (Fig. S1F). GO analysis of 1,281 upregulated luminal DEG genes (Fig. 2A) primarily showed enrichment of pathways related to metabolism and catabolism of lipids (Fig. S1E).

**Figure 2:**
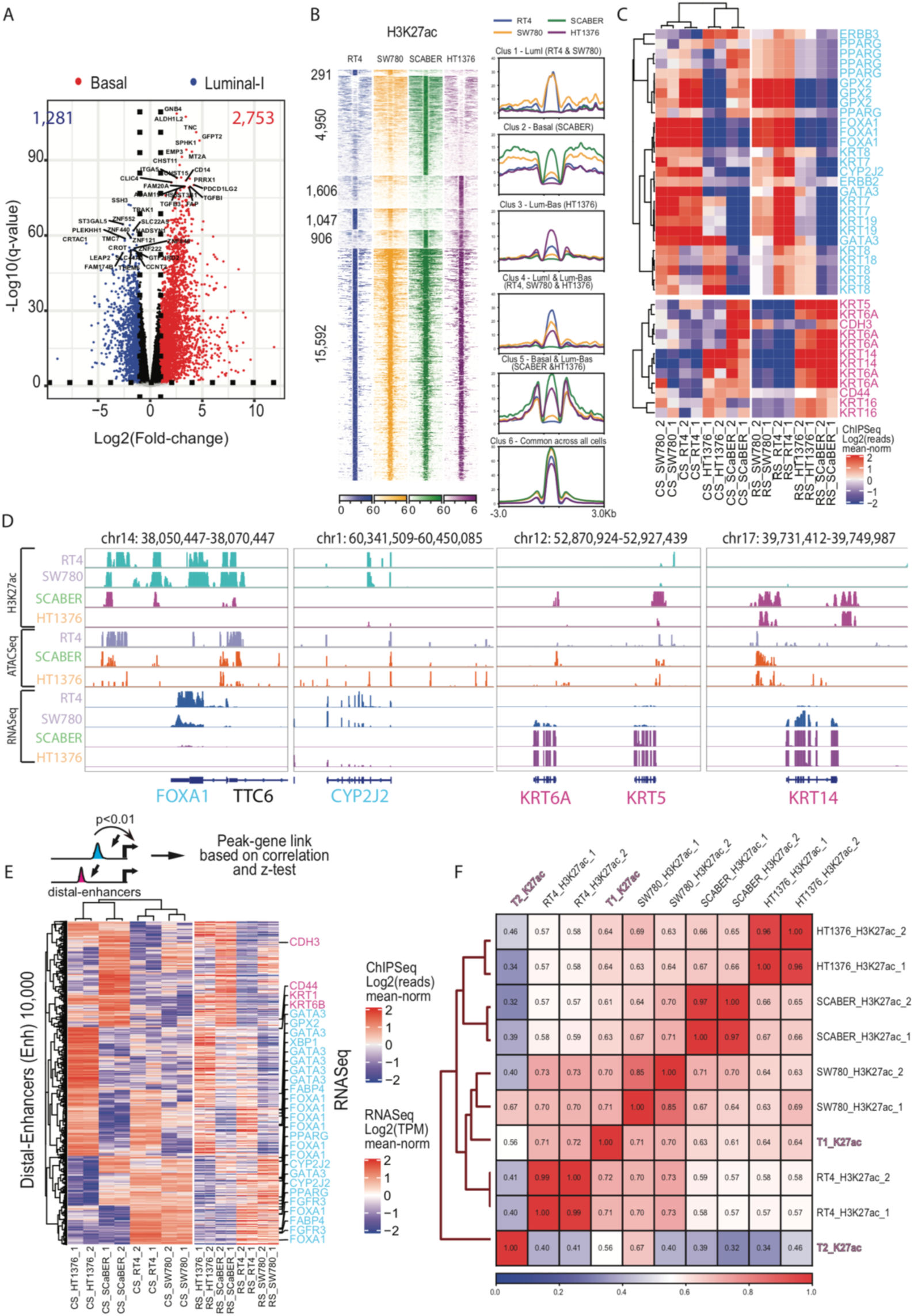
Luminal and basal bladder cancer subtypes are regulated by distinct epigenetic promoter and distal -enhancers. A) Differential expression gene (DEG) analysis of Luminal I and Basal cluster shows 2,753 basal-specific upregulated genes and 1,281 luminal I specific upregulated genes. B) A comprehensive distinct set of promoter H3K27ac ChIP-Seq tags for six selected clusters are shown here genome-wide (left). Signal H3K27ac intensity profiles for each BLCA cell line is plotted for each identified cluster indicating its specificity (right). C) Corresponding promoter H3K27ac and its associated RNA-Seq signals for selected luminal and basal genes shows remarkable similarity. D) Genome browser signal tracks for luminal genes FOXA1, CYP2J2 and basal genes KRT6A, KRT5 and KRT14 incorporating H3K27ac, ATAC-seq and RNA-seq data collected from RT4, SW780, SCaBER and HT1376 are shown. E) Integrated H3K27ac peaks at distal-enhancers and RNA-Seq gene expression association model identifies putative enhancers and gene regulation. Top 10,000 most variable enhancers (left heatmap) are plotted along with their corresponding gene expression (right heatmap). Luminal (cyan) and basal (magenta) genes are highlighted for their specific linked enhancers. F) Genome-wide H3K27ac signals for cell lines is correlated with the tumors T1 and T2 to demonstrate similarity of enhancer landscape.

### Luminal and basal BLCA subtypes are regulated by distinct epigenetic promoter and distal enhancers

Enrichment of H3K27ac signals has been used to predict both active promoters and distal enhancers [28, 29]. Therefore, we first performed ChIP-Seq for H3K27ac in all four cell types and two patient samples. We observed that biological replicates are always clustered first, indicating our results are highly reproducible [Supplementary Table 1]. Further, we found that two luminal subtypes (RT4 and SW780) clustered together, while two basal-squamous (SCABER and HT1376) cell lines are grouped together as well (Fig. S2A). The clustering result also suggests that global epigenomic profiles accurately reflects cell identity. The hierarchical clustering in the cell lines based on H3K27ac signals was also mirrored by the global mRNA expression by RNA-Seq data (Fig. S2-B, S2C).

Next, we examined promoter usage based on H3K27ac signals at known genes. We confirmed that promoter H3K27ac intensities are remarkably similar to gene expression (Fig. 2B), and the clustering analysis based on promoter H3K27ac intensity still distinguishes luminal and basal BLCA models (Fig. S2-C). For example, we observed that two luminal subtype BLCA cell lines RT4 and SW780 have similar H3K27ac patterns at luminal genes *FOXA1, GATA3* and *PPARG* (Fig. 2C and Fig. 2D). HT1376 is marked as luminal-basal subtype, and it accordingly shows a similar promoter H3K27ac pattern at luminal genes (*GATA3, ERBB2/3, KRT7/8/18/19*) with other luminal cells (RT4 and SW780; Fig. 2C and Fig. 2D), but also shows enrichment at basal genes (*KRT6A, KRT14* and *KRT16*), similar to the basal-squamous cell line SCABER.

Distal H3K27ac peaks from gene promoter regions have been used as markers for active enhancers [28, 30]. We took the same approach here and on average, we predicted 59,466 (40,731 – 78,506) enhancers in each cell line. To link the distal enhancers to their target genes, we performed a correlation-based distal-enhancer peak-gene association as described by Greenleaf 2018 [31], and identified the top 10,000 variable distal-enhancers that show significant correlation to its linked gene (correlation ≥ 0.5, p<0.01; a total of 58,509 satisfied our criteria Fig. 2E). We observed that enhancer clustering pattern was similar to the RNA-Seq patterns where all the enhancers linked to known luminal genes (such as *FOXA1, GATA3, PPARG, CYP2J2, FGFR3*, etc.) were predominantly shared between luminal cells (RT4 and SW780) and luminal-basal cell line HT1376 (Fig. 2E and Fig. S2-D). This suggests that regulation of the aggressive basal BLCA subtype may require epigenetic activation of its promoter’s histone modification at known basal genes (such as *KRT14* and *KRT6*) and as well as loss of distal enhancers linked to luminal genes. Moreover, to understand the clinical relevance of our findings, we performed H3K27ac ChIP-Seq in two muscle invasive bladder patient samples. Our results show a remarkable correlation of tumor T1 with luminal subtype of bladder cancers (Fig. 2F) while tumor T2 belonged to a distinct cluster possibly due to low number of enriched peaks detected in T2 (28,497) compared to T1 (166,591). Our results show that the epigenetic regulation is a key contributor to molecular subtype assignment in both commonly used models of BLCA and in patient-tumors.

### Distinct set of transcription factors in luminal and basal BLCA regulatory elements

We performed ATAC-Seq in RT4, SW780, SCABER and HT1376 cell lines to evaluate their open-chromatin status in the genome. On average, in each cell, we identified 28,671 open chromatin regions (Fig. 3A and Supplementary table S3). Among them, 43.7% at promoter region and 56.3% at distal regions. Overall, > 90% of the open chromatin region at promoter overlap with H3K27ac (Fig. S3-A). The overlap of distal ATAC-Seq peaks and H3K27ac are lower (Fig. S3-A). Genome-wide correlation of ATAC-Seq showed that the HT1376 and SCABER clustered together with 80% similarity (Fig. S3-C) compared to luminal RT4 (∼65%). We noted that this observation agrees with the RNA-Seq based clustering (Fig. S2-B) and H3K27ac based clustering (Fig. S2-A).

**Figure 3:**
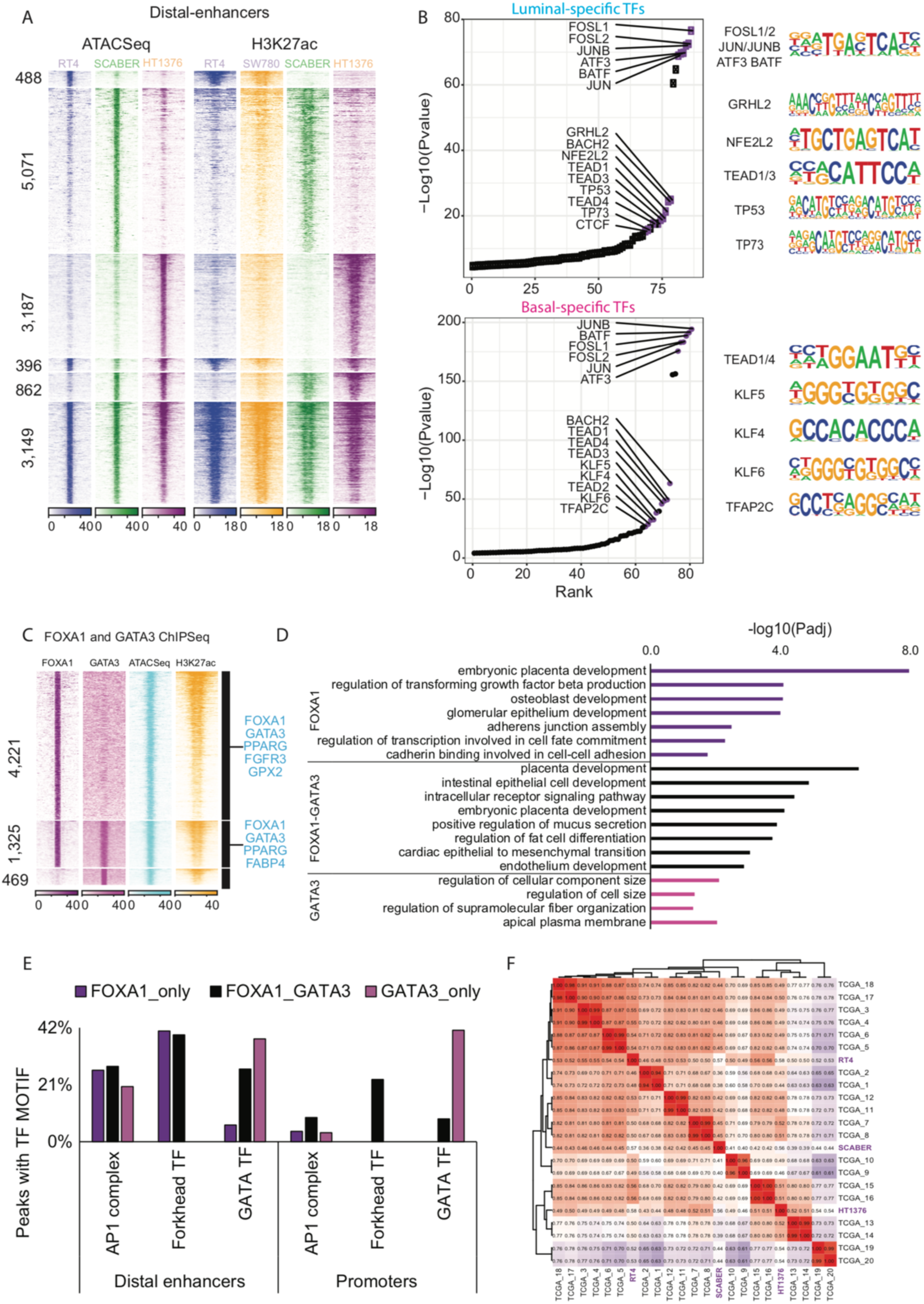
Distinct set of transcription factors in luminal and basal bladder cancers regulatory elements. A) A comprehensive and a distinct set of open-chromatin ATAC-Seq signal tags located at previously identified six enhancer clusters are shown here genome-wide B) TF motif analysis results is shown here as a ranked plot (left) and motifs (right), where for luminal-specific (top) and basal-squamous-specific open-chromatin enhancers (bottom). C) FOXA1 and GATA3 bound open chromatins located at distal-enhancers of RT4/luminal cell line is depicted here in three groups: FOXA1 only, GATA3 only and FOXA1 and GATA3 bound sites. D) Gene ontology analysis of pathways for each group of binding sites (FOXA1 only, FOXA1 and GATA3 and GATA3 only) is shown. E) Observed occurrence of TF motifs (AP-1, FOX Forkhead and GATA) is shown here at distal-enhancers and promoters of three groups. F) Genome-wide open chromatins of four BLCA cell lines show similarity in TCGA bladder tumors (Corces, Granja et al. 2018).

Next, we performed motif analysis of these open-chromatin regions. We observed that CTCF and AP-1 complex are enriched in all cell lines (Fig. 3B and Fig. S3-F). Further ranking of TF motifs by its enrichment p-value showed that luminal open chromatins (shared between RT4, SW780 and HT1376) were enriched for GRHL2, TP53 and TP63 binding motifs while basal open-chromatins (shared between SCABER and HT1376) were enriched for TEAD1/4, KLF factors and TFAP2C (Fig. 3B) binding motifs. GRHL2 [32] and TFAP2C [16] were previously reported to be a luminal and basal-squamous-specific genes, respectively, thereby validating our findings. Interestingly, binding motifs for AP-1 complex proteins FOSL1/2 JUN/JUNB, ATF3 and BATF TFs [33] were the topmost enriched motifs for both luminal and basal-squamous open chromatins. We then comprehensively mapped all the enriched TF motifs in luminal, basal-squamous and shared open chromatins of distal enhancers to discover the relationship between TFs and BLCA subtypes (Fig. S3-D). We discovered that at distal enhancers, the luminal BLCA subtypes are uniquely driven by previously reported steroid hormone receptor TFs including PPARG [4, 6, 8, 16] and orphan nuclear receptors (ESSRA/ESSRB) [4, 34], as well as thyroid hormone receptor (THRB), which has not been previously implicated. On the other hand, basal-squamous open chromatin areas at enhancers show unique enrichment of previously unreported factors MADS box TF MEF2C and the homeobox TF OTX2. Not surprisingly, luminal pioneering TFs such as forkhead transcription factors (FOXA1/2/3, FOXF1, FOXK1, FOXM1), and GATA TFs (GATA3/4/6) were enriched in luminal-specific enhancers with an open chromatin conformation. More surprisingly, forkhead and GATA motifs were also identified as being associated with open chromatin at enhancer elements across cell lines (Fig. S3-D). While FOXA1 is known to have low expression in basal bladder cancers cell lines and tumors (Fig. S1-A, S1-B and S2-B), the enrichment of forkhead and GATA motifs in open chromatins of all BLCA cell lines suggests a more general role for other members are these large families of transcriptional regulators. In addition, this may indicate that luminal-specific TFs such as FOXA1 and/or GATA3 may be poised to bind to these areas of open chromatin. Additionally, FOXA1 and GATA3 are known to play a role in the development of urothelium [32] suggesting that their binding sites may be primed early during development. We also discovered that the stem-cell associated pioneering TFs such as KLF factors (KLF10/14) ATF factors (ATF1/2/4/7) and NANOG were enriched in basal bladder enhancers. This is interesting because there exists a progenitor cell population within basal urothelium that can contribute to urothelial development and differentiation [35, 36].

### FOXA1 and GATA3 bind at luminal open-chromatins at regulatory distal enhancers to drive expression of luminal-specific genes

We hypothesized that the TFs such as FOXA1 and GATA3 binds at the open chromatin region to pioneer luminal enhancers and activate associated gene expression. To test this hypothesis, we performed GATA3 ChIP-Seq in the RT4 luminal BLCA cell line and obtained FOXA1 ChIP-Seq in RT4 cells from our previously published work [8]. As predicted, luminal TFs FOXA1 and GATA3 showed enriched binding at the open chromatin loci of luminal (*FOXA1, GATA3, PPARG, FGFR3* and *FABP4*) distal enhancers (Fig. 3C). More specifically, we discovered that 1,325 distal enhancers that show co-binding of both FOXA1 and GATA3 in RT4 (Fig. 3C). Similarly, FOXA1 and GATA3 showed enriched binding at open chromatin loci of luminal (*FOXA1, ERBB3, KRT19, GPX2* and *FABP4*) promoters (Fig. S3-E).

GO term analysis of genes proximal to these distal enhancer sites showed regulation of TGF beta production, epithelium development, regulation of transcription involved in cell fate commitment and cell-cell adhesion biological processes (cadherin binding and adherens junction assembly) as terms associated with FOXA1. In addition, regulation of cellular component and cell size and apical plasma membrane biological processes were terms identified with GATA3-bound genes proximal to these distal enhancers, suggesting a strong involvement of both TFs in committing to cell-fate and luminal differentiation (Fig. 3D). In regard to proximal genes associated with distal enhancers bound by both FOXA1 and GATA3, terms identified were associated with various developmental processes and the regulation of mucus secretion and fat cell differentiation, important metabolic attributes of differentiated urothelium (Fig. 3D).

We then proceeded with the motif analysis of FOXA1 only, GATA3 only and co-bound sites. Surprisingly, AP1-complex was enriched specifically in all distal enhancers in addition to FOXA or GATA motifs (Fig. 3E). The order of binding of these three factors remains to be investigated. Finally, to understand the clinical relevance of our findings, we compared our four BLCA cell lines to the TCGA muscle-invasive bladder tumor ATAC-Seq data [31] and discovered that the genome-wide open-chromatin profile in our cell lines are clustered with distinct clusters of tumors (Fig. 3F), suggesting that the open chromatin regions in these cell lines share similar patterns with patient tumors.

### Luminal and basal subtype of BLCA show distinct 3D genome organization

Previous studies have shown that 3D chromatin organization is associated with epigenetic activation or silencing of genes in cells [37]. For example, the majority of heterochromatin is known to be compressed in nuclei and located near the lamina associated periphery of the nuclear envelope [37]. To obtain a genome-wide 3D landscape of luminal and basal BLCA, we performed high-resolution Hi-C experiments on all four cell lines and two bladder tumor patients (Fig. S4). Each sample was sequenced for at least 300M reads each. We used our recently developed software, Peakachu [38], which is a machine learning based chromatin loop detection approach, to predict loops at 10Kb bin resolution. First, we identified 58,988 and 49,258 loops in SCABER and SW780 cells respectively (prob>0.8). Then by using the probability score output from Peakachu, we defined subtype-specific chromatin loops, as shown in the Aggregate Peak Analysis (APA, Fig. 4A) [39]. While most loops (42,949) are shared between the basal and luminal cell lines, we observed more basal-specific loops in SCaBER (16,039) relative to luminal-specific in SW780 (6,309). We then compared each of these categories with loops detected in 2 patient samples (Fig. 4B): ∼70% of common 3D chromatin loops identified in both SW780 and SCABER were validated in either tumors (T1 or T2) or both (T1 and T2). Consistent with previous H3K27ac ChIP-Seq data, which showed that both T1 and T2 were like luminal BLCA cell lines RT4 and SW780, more than 50% luminal specific loops identified in SW780 were present in the T1 or T2 loop set, while only ∼40% of basal specific loops could be validated by patient Hi-C data.

**Figure 4:**
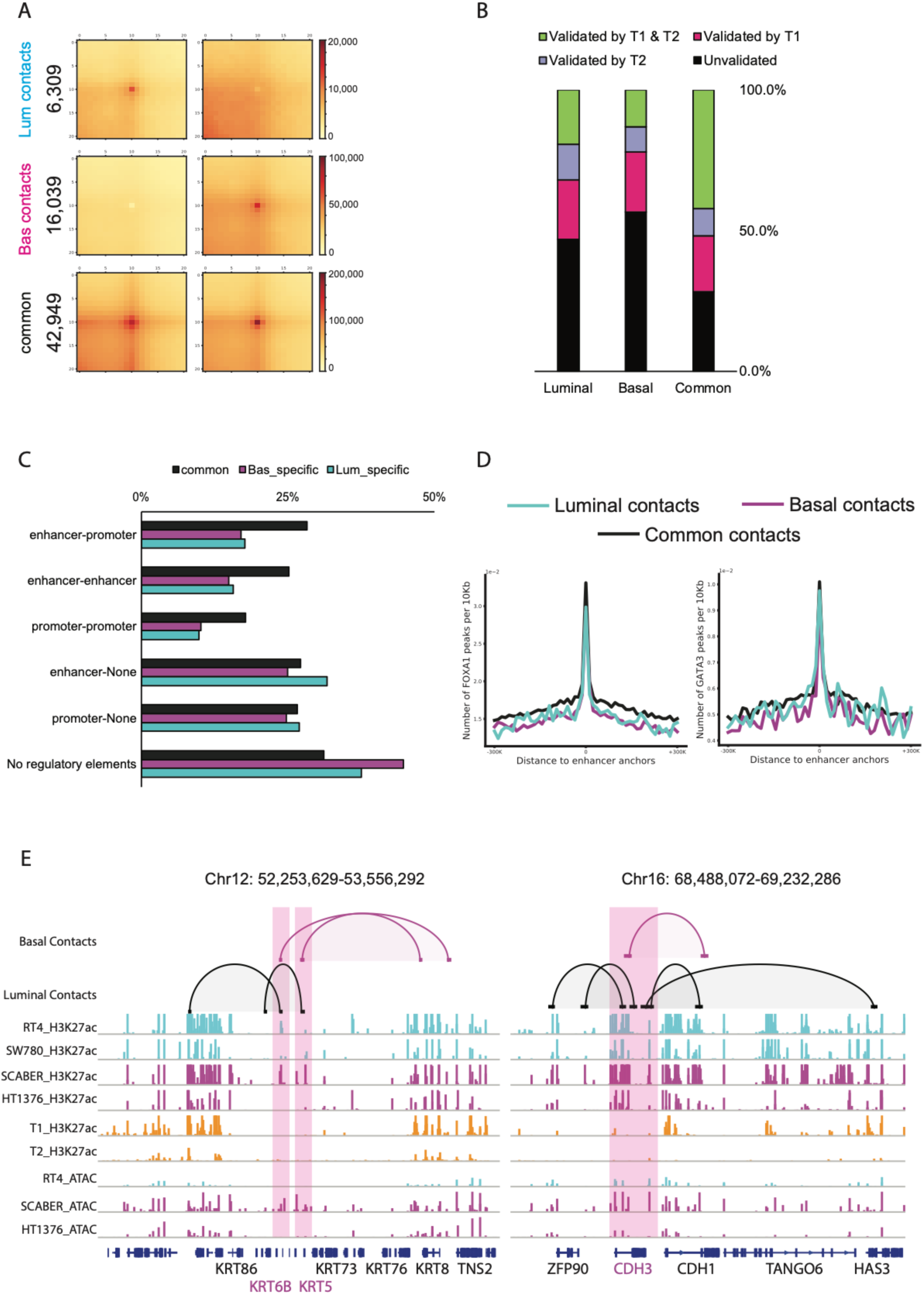
Luminal and basal subtype of bladder cancers show distinct 3D genome organization. A) Hi-C loop analysis of luminal and basal-squamous cell lines show distinct luminal contacts, basal-squamous contacts and commonly shared contacts. B) Contacts identified in luminal and basal-squamous cell lines are shared and validated in muscle-invasive bladder tumor T1 and T2. C) The type of contact based on the overlap of contact location at either enhancer (H3K27ac at distal region) or promoter (H3K27ac and H3K4me3 at promoter) is shown. D) Enrichment of FOXA1 (left) and GATA3 (right) peaks is shown here at loop anchors of either luminal, basal or common contacts. E) Selected basal genes (KRT6B, KRT5 and CDH3) that contain basal and common enhancer-promoter loops are shown here as genome browser tracks.

Next, we interrogated the roles of different epigenetic regulatory features within the different loop category. We first defined promoter and enhancer elements as regions that overlapped with either H3K4me3 or H3K27ac ChIP-Seq peaks. Among the 42,929 shared loops in both cell lines, 28% contain enhancers in one anchor and promoters in another, 25% contain enhancers in both ends, and 31% contain no defined elements at either anchor (Fig. 4C). For basal or luminal specific contacts, although the enrichment for enhancers and promoters is overall decreased relative to common contacts, we found a similar or even higher ratio of contacts containing enhancers at a single anchor, suggesting specific regulatory events might exist in each bladder cancer subtype. Furthermore, we found a significant FOXA1 and GATA3 enrichment at these enhancer contact anchors, indicating the involvement of these pioneer factors in the regulation of the 3D genome (Fig. 4D). This finding is in agreement with previous studies reporting the enrichment of FOXA1 binding sites in enhancer-promoter contacts [40].

Finally, we examined enhancer and promoter contacts in each category for driving subtype-specific gene expression. First, we found several common enhancer and promoter contacts in both luminal and basal genes suggesting that several enhancer and promoter contacts were either poised or pre-existing in the absence of TF binding and regulation of its associated gene expression [41]. Second, we found the enrichment of luminal genes (*ERBB2, FOXA1, GATA3, KRT20* and *PPARG*) in luminal contacts and basal genes (*CD44, CDH3, KRT5, KRT6B*) in basal contacts. This suggests that the development of aggressive basal BLCA may require concordant changes to enhancer and promoter 3D looping in addition to the loss of FOXA1/GATA3/AP-1 driven distal enhancers and promoters.

### Neuronal PAS Domain Protein 2 (NPAS2) is a novel luminal BLCA TF which regulates luminal gene expression and cell migration

Genome-wide open chromatin analysis of luminal (shared between RT4, SW780 and HT1376) and basal-squamous (shared between SCABER and HT1376) BLCA cell lines provides an ideal platform for the identification of novel transcriptional regulators of BLCA cell fate and phenotype. Our approach identified the enrichment of several TF motifs with a common basic helix-loop-helix (bHLH) signature such as MYC, MYCN and MAX amongst others (Fig. 5A). Since bHLH TFs such as MYC are bona fide oncogenes, we systematically mapped the mutually exclusive segments of bHLH TFs that appear at luminal-specific, basal-specific and shared areas of open chromatin (Fig. 5A – Venn diagram). This approach identified several TFs such as ARNTL, HIF1A, TFE3, MYC and MAX which exhibited patterns of expression similar to genome-wide signatures in TCGA basal-squamous tumors. This approach additionally identified a set of TFs, including MYCN, REPIN1, BHLHE41 and NPAS2 which exhibited patterns of expression similar to luminal tumors (Fig. 5B). GO term analysis of these luminal bHLH TFs revealed that they play a major role in circadian regulation of genes (NPAS2, ARNTL, BHLHE41, MYC, HIF1A; Fig. 5C). Analysis of oncologic outcomes included in the TCGA dataset [27] identified the novel bHLH containing regulators NPAS2 (Fig. 5D) and BHLHE41 (data not shown) in luminal bladder cancers whose expression is significantly correlated to overall patient survival.

**Figure 5:**
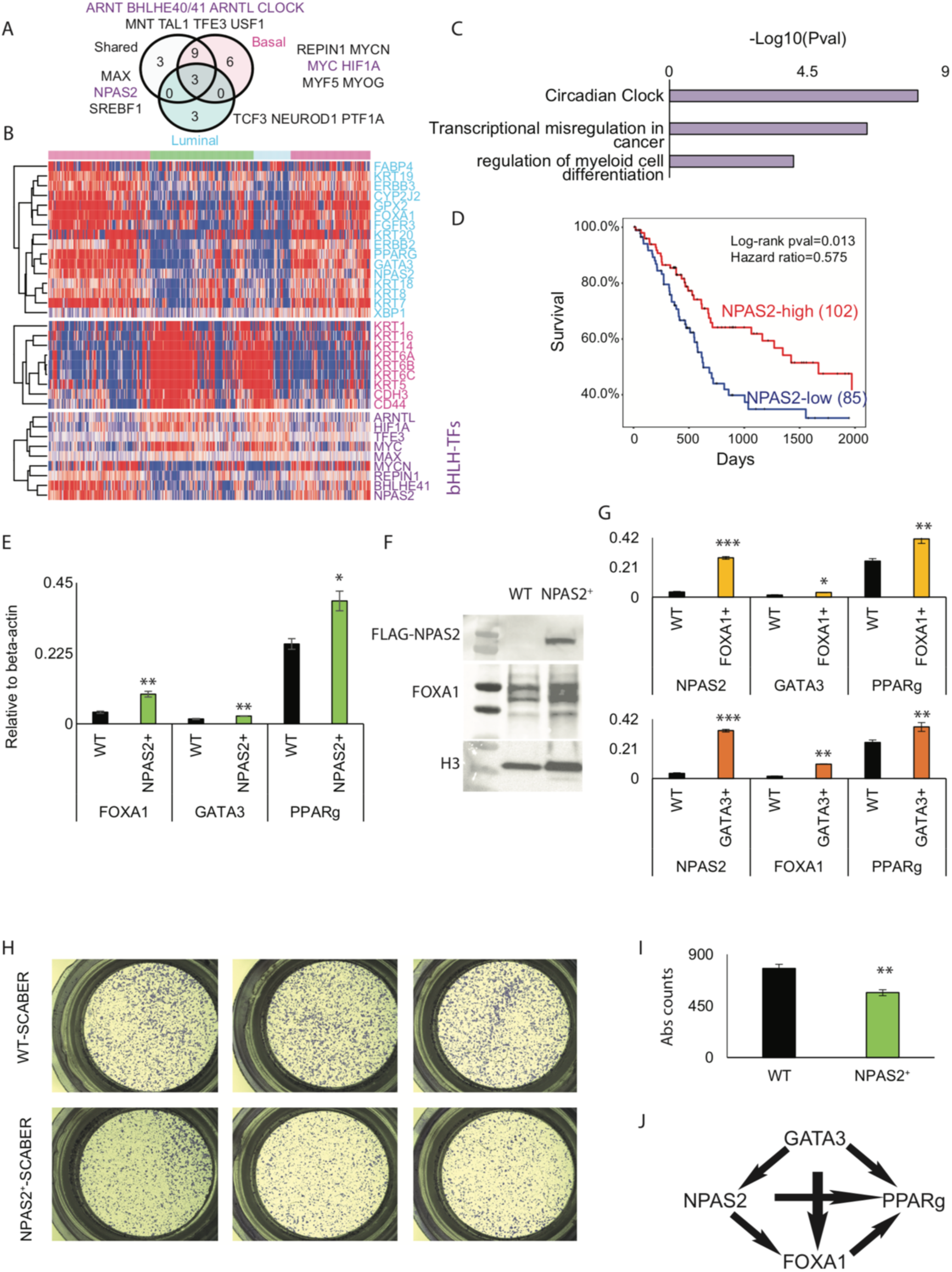
NPAS2 is a novel luminal bladder cancer TF which regulates luminal gene expression and cell migration. A) Several basic helix-loop-helix (bHLH) TFs are prevalent in open-chromatin regions of luminal, basal and common (shared) BLCA cells. B) Hierarchial clustering of TCGA’s tumor RNASeq reveals TF expression pattern is concordant with BLCA molecular subtype. Top heatmap is luminal, whereas middle and bottom heatmap include signature genes for basal-squamous BLCA and bHLH TF genes, respectively. C) Expressed TFs were computed for gene ontology analysis and –Log10(pvalue) for enriched terms is depicted. D) NPAS2 Kaplan Meier curve is shown here for 2,000 days with log-rank statistics and hazards ratio. E) RT-qPCR results for FOXA1, GATA3 and PPARG genes are shown here for control and NPAS2 overexpressed SCABER basal cell line. F) Western blotting analysis for FLAG-NPAS2 and FOXA1 following FLAG-NPAS2 overexpression in the basal-squamous cell line SCaBER. G) Luminal gene RT-qPCR is shown here for control and FOXA1/GATA3 overexpressed SCABER basal cell line. H-I) Transwell migration assay representative crystal violet staining (H) and (I) quantification of differences in transwell migration are shown following overexpression of NPAS2 in SCaBER. J) NPAS2 regulation of luminal genes is shown here as a model.

To determine if NPAS2 expression influences the expression of luminal-specific genes, we overexpressed NPAS2 in the basal-squamous BLCA cells line SCABER and performed PCR and western blotting analysis. Overexpression of NPAS2 in SCABER resulted in significantly increased expression of *FOXA1* and *PPARG*, with a limited impact on the expression of *GATA3* (Fig 5E). Western blotting analysis confirmed increased expression of FOXA1 at the protein level (Fig. 5F). Because our functional genomics analysis suggests FOXA1 and GATA3 cooperate to regulate luminal target genes, we individually overexpressed FOXA1 and GATA3 in SCABER cells to test their ability to regulate NPAS2 expression. We observed increased expression of NPAS2 by both FOXA1 GATA3 (Fig. 5G). To additionally test the impact of NPAS2 on oncogenic behavior, we performed trans-well migration assays. Overexpression of NPAS2 in SCABER decreased trans-well migration (Fig. 5H and 5I). Since basal BLCA are frequently identified at a relatively advanced stage, our observation suggests that low expression of NPAS2 promotes malignant behavior. Based on these observations, we propose a model in which NPAS2 regulates luminal gene expression by controlling expression of key TFs (Fig. 5J).

## Discussion

Muscle invasive BLCA is a morbid and expensive disease to treat [3, 42-44]. The high prevalence of early stage disease in newly diagnosed patients, results in significant financial burden associated with monitoring, administering and managing care. However, with recent development of immunotherapies such as anti-PD-1 [45] and PD-L1 [46], as well as targeted approaches including FGFR3 inhibitors has revolutionized clinical management. [47]. However, response rates to these and other standard approaches are suboptimal, suggesting the need for an increased molecular understanding. In keeping with this, recent National Comprehensive Cancer Network (NCCN) guidelines have encouraged biomarker and molecular-based subtype studies to further stratify patients for recent targeted therapies [48].

It has been suggested that RNA-Seq based molecular subtyping of BLCA are prognostic of clinical outcomes in patients [6-8]. While TCGA and other studies have identified mRNA based molecular subtypes, the epigenetic differences underlying these expression subtypes is unknown. The Encyclopedia of DNA Elements (ENCODE) consortium has contributed greatly to the current understanding of how epigenetic modifications in multiple tissues vary to regulate tissue-specific gene expression [30]. Histone modification states such as H3K27ac amongst various other epigenetic states mark enhancers and promoters that form an complex interacting network hub to regulate gene expression [49, 50]. TCGA has incorporated DNA methylation to understanding epigenetic states [27]. DNA methylation states have been shown to be coupled with histone modifications particularly at the CpG cites at promoters [51]. However large changes in epigenetic histone modification states that influence gene expression lie in distal regions in enhancers and other sites that orchestrate the 3D genome (CTCF) [30, 40, 52-54]. Hence, our study utilized large-scale genomic experiments such as ATAC-Seq, H3K27ac ChIP-Seq, as well as FOXA1 and GATA3 ChIP-Seq and Hi-C combined with RNA-Seq to construct a deep molecular map of luminal and basal BLCA in both cell line models as well as patient tumors. We have further utilized CCLE and TCGA datasets to orthogonally validate and derive inferences for clinical importance of our findings.

We found evidence for regulation of luminal and basal bladder cancer genes by proximal-promoters and distal-enhancers that form long-range chromatin contacts and potentially drive oncogenic programs. Our findings were in large agreement to previously known work on the role of FOXA1 and GATA3 in the literature in regulation, maintenance and oncogenic programs of luminal bladder cancers [8, 19-21]. Interestingly, we found a novel co-regulation of FOXA1 and GATA3 with a common binding partner of AP-1 complex like breast cancers [55, 56] and drive distal-enhancers, but not promoters. Our comprehensive 3D genome map shows distinct chromatin contact interaction networks available to luminal and basal BLCA. To our knowledge, this is the first report of a comprehensive 3D genome map of bladder tumor patients. Our analysis further demonstrates the regulation of distal-enhancers with promoters of BLCA subtype-specific genes through physical contacts as reported in other studies [39, 40, 50]. Through our analysis, we have identified novel bHLH TF regulators of luminal BLCA – NPAS2 whose expression correlates with overall survival of BLCA patients included in the TCGA cohort. Through several biological experiments we showed that NPAS2 intricately regulates luminal TFs *FOXA1, GATA3* and *PPARG* gene expression, and further diminishes the migration ability of basal BLCA. Most importantly, our work highlights how these TFs can cooperatively regulate molecular subtypes and drive clinical associations. One limitation of our study however was the lack of a consortium-level analysis of ChIP-Seqs for histone modifications and all major TFs. Such an approach would increase precision in the regulatory analysis. Additionally, the availability and cost of sequencing limited our current capability to include a large set of patient tumors. However, previous studies were limited to the understanding of single or combination of few TFs in the context of gene regulation. Therefore, we believe that our study will emerge to be a solid comprehensive resource to launch a further series of hypothesis-driven biological experiments based on gene and epigenetic perturbations to unveil both novel molecular targets as well as biomarkers.

## Acknowledgements

F.Y. is supported by 1R35GM124820, R01HG009906, U01CA200060 and R24DK106766.

## Author Contributions

The study was conceived and supervised by F.Y. and D.D.. T.J., B.Z., Q.J. H.Y., T.L, J.X., and H.Y. performed the experiments. T.J., X.W., F.S., and Y.L. performed data analysis. T.J. and B.Z. performed ATAC-Seq, ChIP-Seq and Hi-C experiments in both cell lines and primary tissues. T.J., B.Z., D.D., and F.Y. wrote the manuscript. All authors approved the final version for publication.

## Materials and Methods

Cell lines and patient tumor samples

Bladder cancer cell lines – RT4, SCABER, SW780 and HT1376 were thawed and cultured as per standard protocol. Bladder tumor samples were obtained from Penn State Hershey, College of Medicine’s biobank storage at the Institute of Personalized Medicine (IPM) with appropriate protocol approval from the institutional review board (IRB). Samples were selected based on its availability (50mg) for several rounds of sequencing experiments.

### Cell culture

Bladder cancer cell lines were cultured in growth medium containing media + 10% fetal bovine serum and 1% penicillin and streptomycin (Corning). RT4, SW780, SCABER and HT1376 cells were cultured in Mcoy’s 5A (Gibco), RPMI-1640 (Corning), Eagle minimum essential medium (MEM; GE life sciences), and Eagle MEM with 1% non-essential amino acids (Corning), respectively. Cells were plated in tissue culture plates (TCPs, Corning) – T-25, T-75 or 15cm dishes to further grow and expand in 5% CO_2_ humidified incubator for several different sequencing experiments. For future storage, cells were preserved in 5% DMSO containing growth medium in the vapor of liquid nitrogen. To passage cells, they were washed with phosphate buffered saline (PBS, Corning) and trypsinized for 5 mins to detach cells from the TCPs. They were further spun down at 200 xG to pellet and washed with PBS for further experiments.

### RNA-Seq

For RNA-Seq, RNA was extracted from frozen cell pellets using RNeasy Mini Kit (Qiagen). Extracted RNA was quantitated using NanoDrop (Thermo Scientific). SureSelect strand-specific RNA library preparation kit (Agilent) was used to generate cDNA libraries where polyA RNA was pulled down using 2 μg of oligo (dT) beads. Extracted cDNA was then fragmentation, reverse transcribed, end repaired, 3′-end adenylated, adaptor ligation and subsequently amplified and bead purified (Beckman Coulter). Barcode sequences were thus used to multiplex high-throughput sequencing. The cDNA library was QC’ed for size distribution and concentration using BioAnalyzer (Agilent) High Sensitivity DNA Kit (Agilent) and Kapa Library Quantification Kit (Kapa Biosystems). Finaly libraries were then pooled and diluted to 2 nM and subsequently sequenced using Illumina HiSeq 2500 (Illumina).

### Chromatin crosslinking and ChIPSeq library preparation

Each cell line was grown in 15cm dishes (x4) with 25mL growth medium were trypsinized as described above and pelleted. Approximately 10-20M cell pellets (2x biological replicates) were crosslinked immediately with 1% formaldehyde in PBS on ice for 5 mins. Crosslinked cells were then washed in PBS and 100μL freshly prepared lysis buffer was added. Lysed cells were then diluted in 900mL TE buffer and sonicated using focused beam ultrasound sonicator (COVARIS). Sonicated samples were repeated for extended periods of time (upto 1.5 hours) until a DNA library size distribution of ∼200-300 bp was achieved. Sonicated DNA-chromatin complexes were then pulled down with anti-H3K27ac antibody and washed several times to remove non-specific interactions. Pulled-down samples as well as direct input controls were all de-crosslinked at 55°C overnight. Samples were treated with RNASe and protein digestion at 37°C and 55°C, respectively, followed by further DNA library extraction using phenol-chloroform method. We then amplified the library using Hi-fidelity KAPA PCR kit (KAPA) for 6 cycles and quantified the library using Qubit high sensitivity DNA assay (Thermofisher). We performed QC using bioanalyzer (Agilent) and then sequenced 20-40M pair-ended reads using Illumina’s HiSeq platform 2500 (Illumina).

### Nuclei extraction and ATACSeq library preparation

Each cell line was pelleted following detachment from the TCPs, as described above. Cells were washed with 500⍰L PBS, counted using hemocytometer and kept on ice as a pellet. We used 50,000 pelleted cells for ATACSeq library preparation as described by Greenleaf [57]. We first extracted the nuclei using chilled and freshly made lysis buffer (TBST) in 500⍰L volume. Then we washed the nuclei pellet again in lysis buffer to remove unwashed mitochondria DNA. We then continued to add the tagmentation DNA (TD) buffer (Nextera) along with Tn5 transposase in 50⍰L volume. We incubated our transposition reaction for 30 mins at 37°C and further extracted the ATACSeq DNA library using PCR purification kit (Qiagen) into 20⍰L elution volume. We then amplified the library using Hi-fidelity KAPA PCR kit (KAPA) for 6 cycles and quantified the library using Qubit high sensitivity DNA assay (Thermofisher). We performed QC using bioanalyzer (Agilent) and then sequenced 40-60M pair-ended reads using Illumina’s HiSeq platform 2500 (Illumina).

### Chromatin crosslinking and HiC library preparation

Each cell line was grown in T-24 plates (x4) with 5mL growth medium were trypsinized as described above and pelleted. Approximately 1-1.5M cells as pellets (2x biological replicates) were crosslinked immediately with 2% formaldehyde in PBS on ice for 5 mins. Crosslinked cells were then washed in PBS and frozen as 1-1.5M aliquots in −80°C for several months before library preparation. We used ARIMA’s Hi-C kit for making libraries (ARIMA Genomics). As per their protocol, we tested and did QC on samples and sequenced 300M-600M reads for each sample based on total usable reads >20Kb.

### Overexpression of NPAS2

To overexpress NPAS2 tagged with FLAG (DYKKK) protein, we obtained plasmids from genscript, expanded the quantity by transforming it in ecoli and picking clones. We then transiently transfected SCABER cell lines with 1⍰g plasmid/well in 6-well plates (2x wells). For transfection we used Lipofectamine 3000 () reagent as per manufacturer protocol and allowed cells to be cultured up to 2 days before collecting it for various analysis.

Reverse transcriptase – quantitative polymerase chain reaction (RT-qPCR)

We used the following primers for RT-qPCR to detect mRNA levels of FOXA1, GATA3, PPARG, NPAS2 and beta-actin.

**NPAS2**: 5’-ACACCCTTTCAAGACCTTGCC-3’ (F); 5’-AGGTTCGTCAACTATGCACATTT-3’ (R)

**FOXA1**: 5’-CGCTTCGCACAGGGCTGGAT-3’ (F); 5’-TGCTGACCGGGACGGAGGAG-3’ (R)

**GATA3**: 5’-ATACACCACCTACCCGCCTAC-3’ (F); 5’-ACTCCCTGCCTTCTGTGCT-3’ (R)

**beta-actin**: 5’-CATGTACGTTGCTATCCAGGC-3’ (F); 5’-CTCCTTAATGTCACGCACGAT-3’ (R)

RNA was extracted from frozen cell pellets using RNeasy Mini Kit (Qiagen). Samples were treated with DNAse to digest any additional DNA extracted during the process. DNAse-free RNA was then further converted to cDNA using reverse transcriptase kit (Takeda) according to manufacturer’s protocol. Reverse transcribed cDNA was then assayed for RT-qPCR using Sybr green PCR kit (KAPA biosystems) at 60°C melting temperature and quantitated using Biorad quantitative PCR system. CT values obtained through the quantitation were then normalized to beta-actin and further transformed to relative expression shown in plots.

### Western Blot

Western blots were performed using standard protocols using BioRad system and digitally imaged using geldoc imager (Biorad) following exposure of the blots to ECL chemiluminescence (Biorad). Briefly, cell lysates were prepared by using RIPA buffer (Invitrogen) in the presence of Complete protease inhibitor cocktail (Sigma). Frozen cell pellets with NPAS2 overexpression or controls were lysed with lysis buffer under ultrasonication pulse at 5% power for 5-10 seconds. Lysates were then pelleted, and supernatants were measured for protein concentration using Bradford assay (Biorad) using plate reader (Biorad). Protein lysates (20-30ug) were then used for running using 10-15% gradient gels (Biorad) for 1.5 hours at 100-110V followed by dry-transfer to PVDF membrane (Biorad dry transfer). Membranes were washed using tris-based saline buffer solutions with 0.1% tween-20 (TBST) and blocking solutions were prepared using 5% w/v western-blot grade dry milk powder (Biorad). Membranes were blocked with blocking buffer for 1 hour and washed between incubations for 5-10 mins (3x). Primary antibody against anti-NPAS2 (Abcam), beta-actin (SantaCruz) were FOXA1 (SantaCruz) diluted at 1:1000, 1:500 and 1:500, respectively, and incubated for 1-2 hours at room temperature. Secondary antibody against anti-Rabbit and anti-Mouse were used at 1:10,000 and 1:5000 respectively, and incubated for upto 1 hour.

### Transwell migration assay

Transwell migration assay was performed using 8⍰m PVDF inserts (Corning) in a transwell chamber fitting into 24 well plates (Corning). Each cells (control and overexpressing NPAS2 for 2 days) were seeded with 50,000 cells in a transwell (x3 replicates) in a FBS free medium containing 1%PS. Cells were allowed to migrate through the transwell inserts for 24 hours into medium containing regular 10% FBS and 1%PS. Transwell chambers were removed, washed with PBS and stained with crystal violet. Cells that were not migrated through the insert were removed using a Q-tip. Migrated cells were then visualized in microscope and scored for number of stained spots and compared with different experiment groups. T-test was used to calculate significance between groups.

---

**Figure S1:**
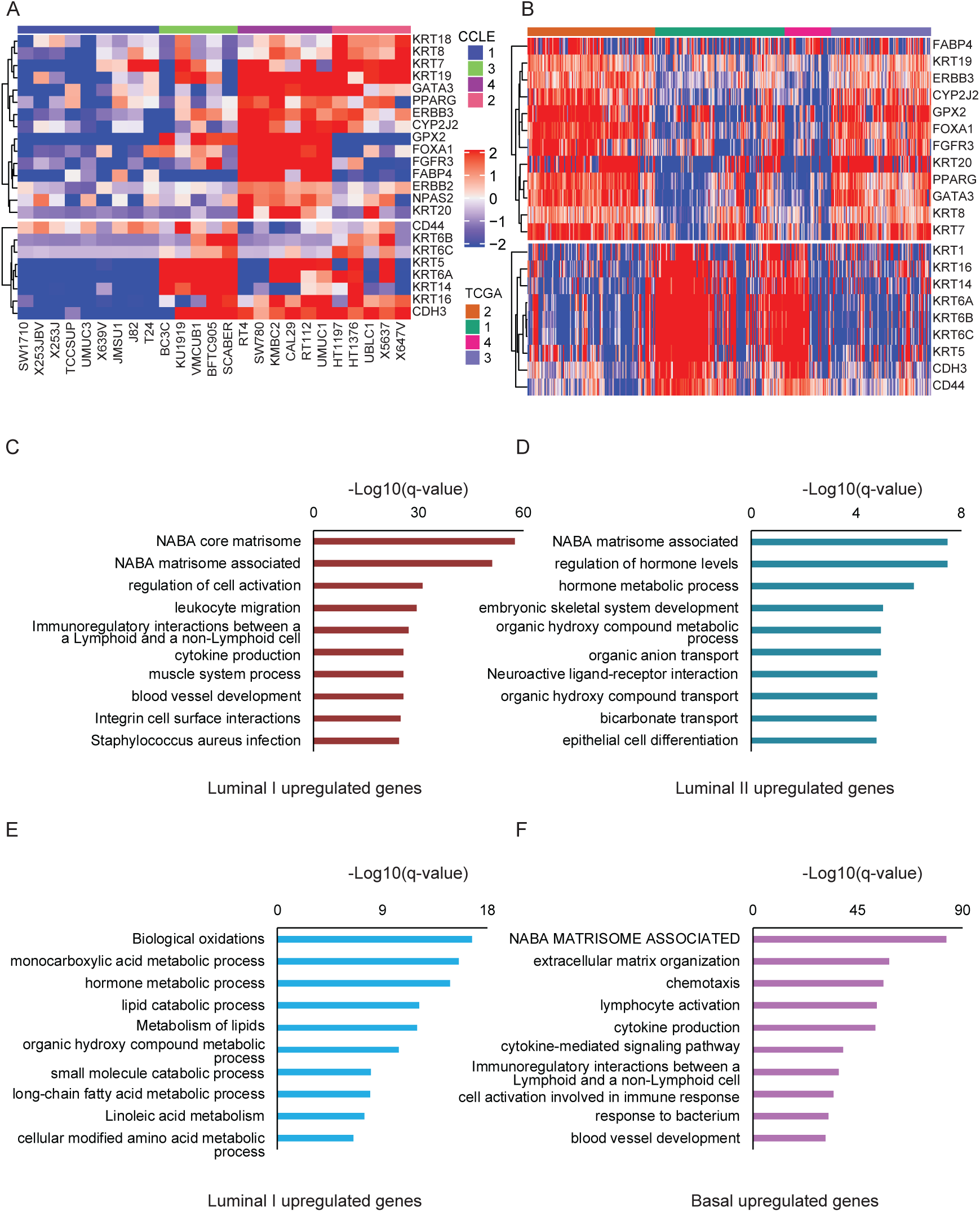
Transcriptomic analysis of luminal and basal subtypes of bladder cancers. A-B) Hierarchical clustering of top 5,000 most variable genes in CCLE (A) and TCGA (B) RNASeq data shows that there are four main clusters of molecular subtypes that show distinct expression for luminal or basal genes. C-D) Gene ontology analysis of Luminal I upregulated genes (C) and Luminal II upregulated genes (D) identified from differential gene expression analysis. E-F) Gene ontology analysis of Luminal I upregulated genes (E) and Basal upregulated genes (F) identified from differential gene expression analysis

**Figure S2:**
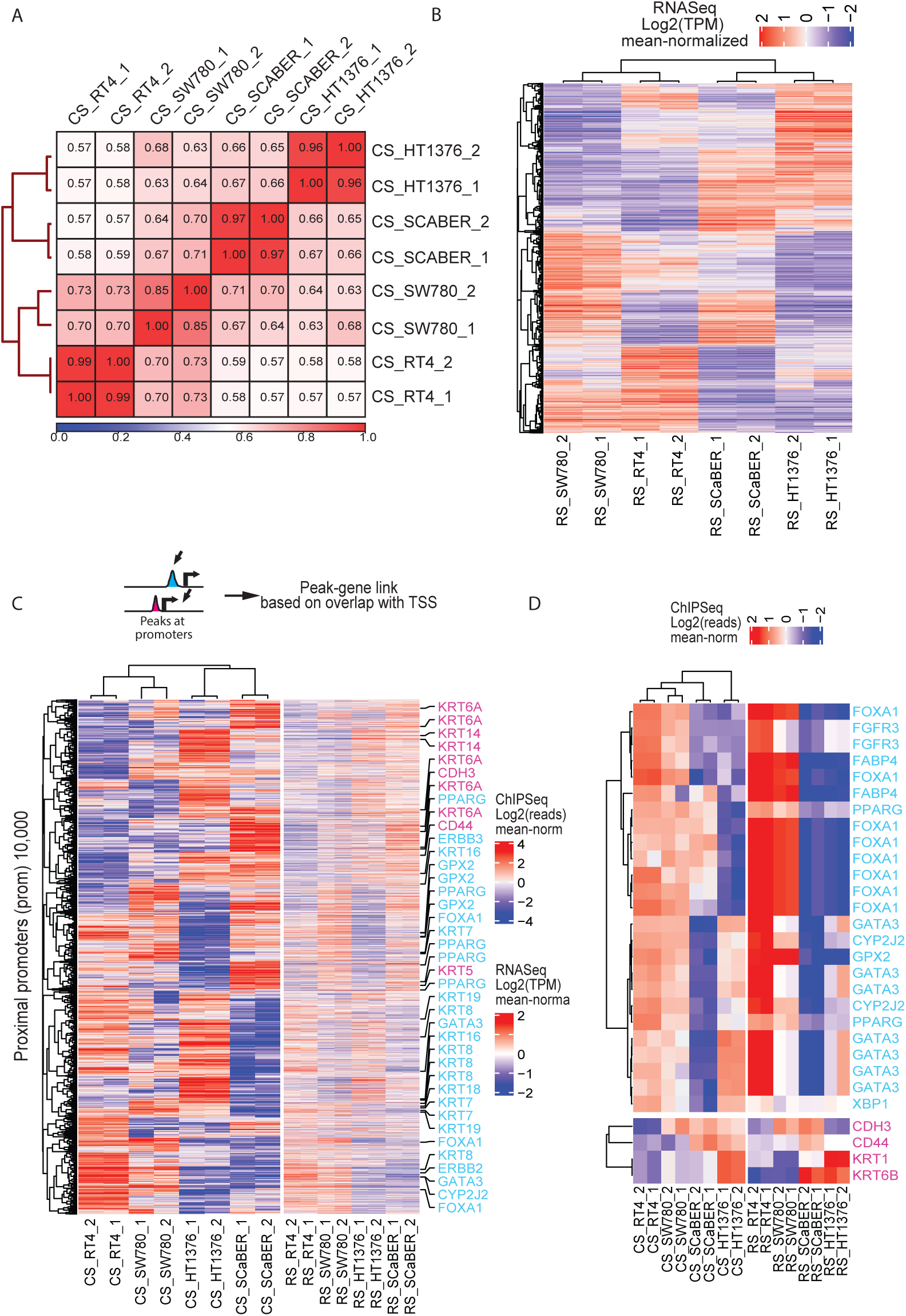
Transcriptomic analysis of luminal and basal subtypes of bladder cancers. A) Genome-wide H3K27ac signals show that biological replicates and as well as molecular subtypes (basal and luminal) cluster together. B) Hierarchical clustering of genome-wide RNA-Seq results for 4 cell lines recapitulate the luminal and basal gene expression based molecular subtypes. C) Integrated H3K27ac peaks at promoters and RNA-Seq gene expression association model identifies putative promoter and gene regulation. Top 10,000 most variable promoters (left heatmap) are plotted along with their corresponding gene expression (right heatmap). Luminal (cyan) and basal (magenta) genes are highlighted for their specific linked enhancers. D) Corresponding enhancer H3K27ac and its linked RNA-Seq signals based on our predicted model for selected luminal and basal genes shows remarkable similarity.

**Figure S3:**
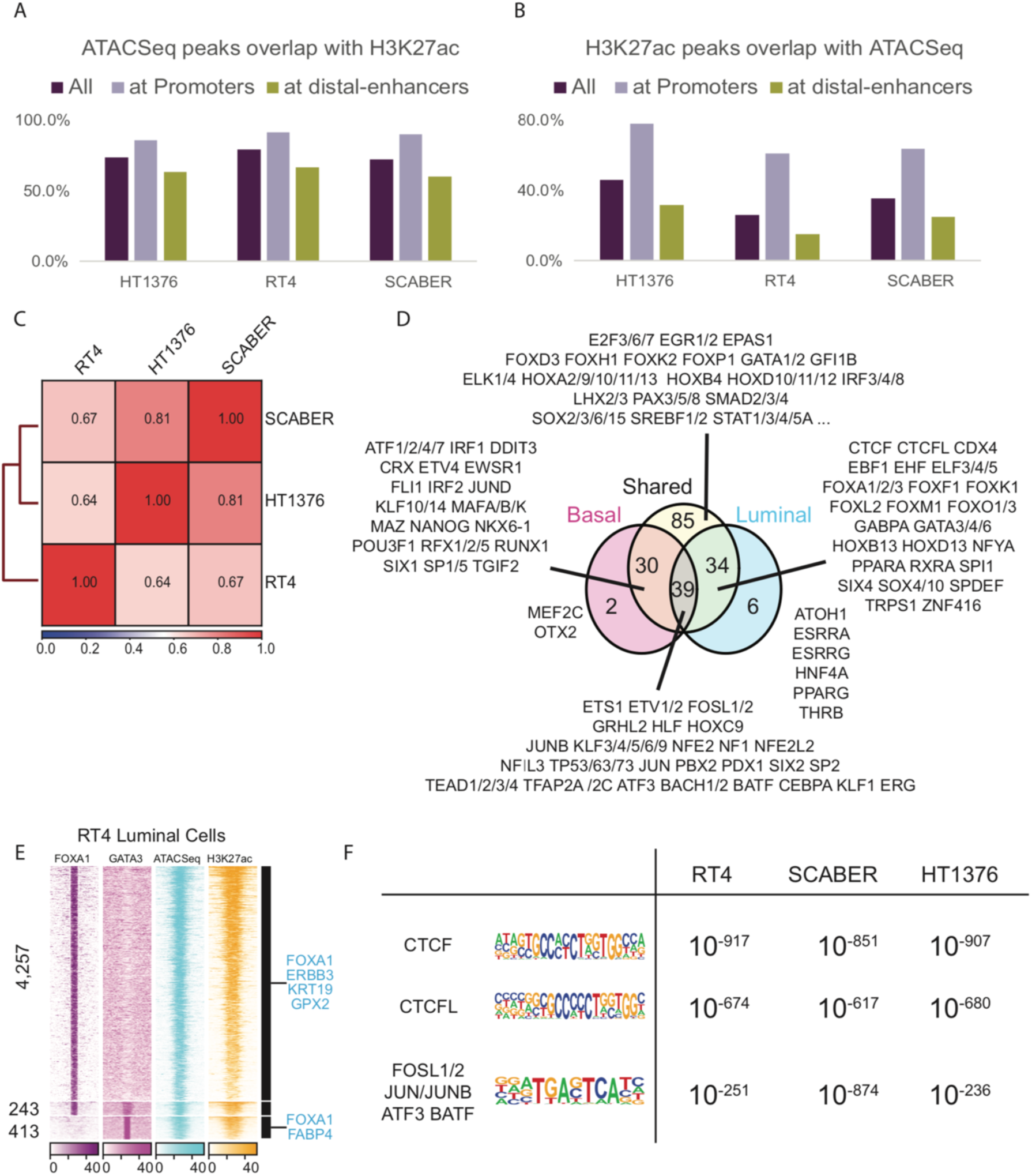
Transcriptomic analysis of luminal and basal subtypes of bladder cancers. A) Genome-wide overlap of ATAC-Seq peaks with H3K27ac ChIP-Seq is shown here for each cell line at either promoter, enhancer or all locations. B) Genome-wide overlap of H3K27ac ChIP-Seq peaks with ATAC-Seq is shown here for each cell line at either promoter, enhancer or all locations. C) Genome-wide correlation of ATAC-Seq signals between cell lines recapitulate enhancer/promoter and RNA-Seq based clustering. D) TFs from motif analysis is mapped out into mutually exclusive and completely exhaustive groups between luminal, basal and common open-chromatins at distal-enhancers. E) FOXA1 and GATA3 ChIP-Seq binding sites overlapped at promoters are shown here as genome-wide tag plot in three groups. F) A comparison of top 3 motifs enriched p-values in each open-chromatins that does not overlap with any H3K27ac signals within its cell lines are shown.

**Figure S4:**
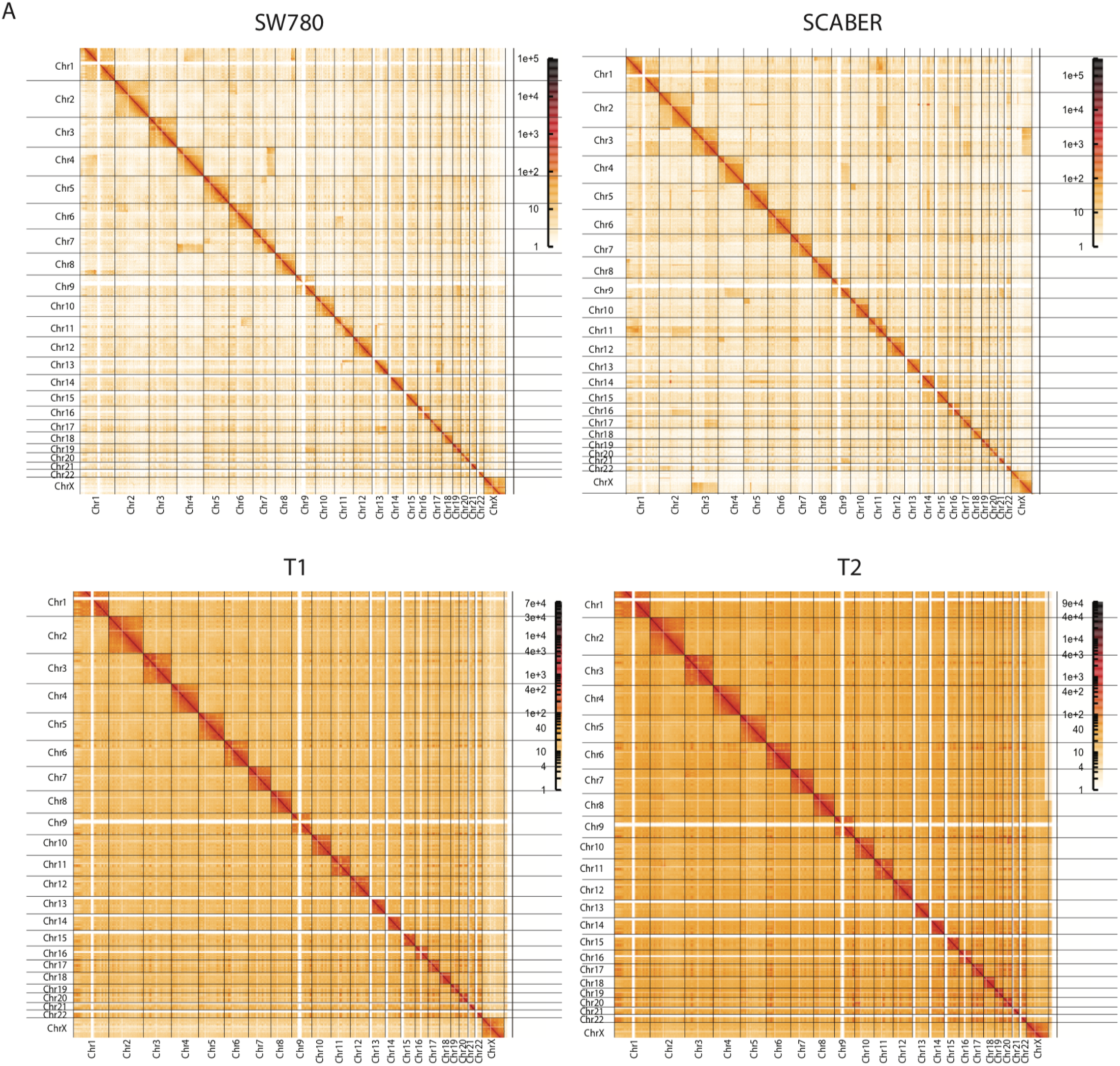
Hi-C maps of luminal and basal subtypes of bladder cancers and bladder tumors. Genome-wide chromosome view of Hi-C map is shown for SW780 (top-left), SCABER (top-right), tumor T1 (bottom-left) and tumor T2 (bottom-right) at 1MB resolution.

**Supplementary Table S1:**
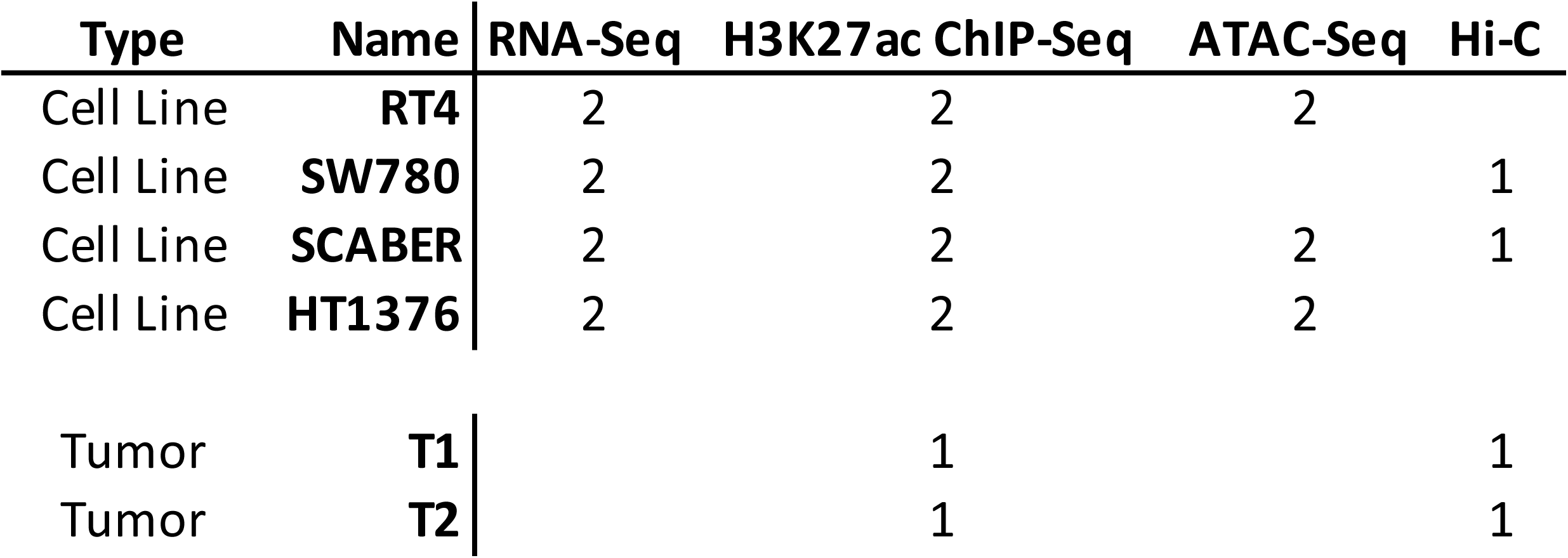
Cell Lines and Experiment table.

**Supplementary Table S2:** RNA-Seq correlation between cell lines.

| **Cell Line** | **Correlation between**<br>**Biological Replicates** |
| --- | --- |
| RT4 | 0.954 |
| SW780 | 0.985 |
| SCABER | 0.966 |
| HT1376 | 0.981 |

**Supplementary Table S3:** ATAC-Seq peaks called at its location.

| **Cell Line** | **Total Peaks** | **Promoters** | **Enhancers** | **%Promoters** | **%Enhancers** |
| --- | --- | --- | --- | --- | --- |
| RT4 | 23265 | 11285 | 11980 | 48.5% | 51.5% |
| SCABER | 33923 | 13089 | 20834 | 38.6% | 61.4% |
| HT1376 | 28827 | 12710 | 16117 | 44.1% | 55.9% |
| **Average** | **28,672** |  |  | **43.7%** | **56.3%** |

